# Two new coral fluorescent proteins of distinct colors for sharp visualization of cell-cycle progression

**DOI:** 10.1101/2020.03.30.015156

**Authors:** Ryoko Ando, Asako Sakaue-Sawano, Keiko Shoda, Atsushi Miyawaki

## Abstract

We cloned and characterized two new coral fluorescent proteins: h2-3 and 1-41. h2-3 formed an obligate dimeric complex and exhibited bright fluorescence. On the other hand, 1-41 formed a highly multimeric complex and exhibited dim red fluorescence. We engineered 1-41 into AzaleaB5, a practically useful red-emitting fluorescent protein for cellular labeling applications. We fused h2-3 and AzaleaB5 to the ubiquitination domains of human Geminin and Cdt1, respectively, to generate a new color variant of Fucci (Fluorescent Ubiquitination-based Cell-Cycle Indicator): Fucci5. We found Fucci5 provided brighter nuclear labeling for monitoring cell cycle progression than the 1^st^ and 2^nd^ generations that used mAG/mKO2 and mVenus/mCherry, respectively.

## Introduction

In recent years, there have been remarkable improvements in our ability to comprehensively unravel the fine details of cellular events. This is owing to the development of the green fluorescent protein from the jellyfish *Aequorea victoria* (avGFP), its spectral variants, such as cyan- and yellow-emitting variants (CFP and YFP, respectively), and GFP-like proteins including red-emitting fluorescent proteins (RFPs) from other organisms. These fluorescent proteins (FPs) can be incorporated into proteins by genetic fusion to develop genetically encoded probes for a variety of cellular functions (Rodriguez et al., 2017).

Fucci is an FP-based probe for visualizing cell-cycle progression (Sakaue-Sawano et al., 2008). The technology harnesses the cell-cycle–dependent proteolysis of Cdt1 and Geminin (Figure 1). Over the course of the cell cycle, SCF^Skp2^ and APC^Cdh1^ E3 ligase activities oscillate reciprocally and the protein levels of their direct substrates oscillate accordingly. Geminin, the inhibitor of Cdt1, is degraded under the control of APC^Cdh1^ E3 ligase. The original Fucci-S/G2/M probe had GFP or YFP fused to the APC^Cdh1^-mediated ubiquitination domain (1–110) of human Geminin (hGem(1/110)); this chimeric protein is the direct substrate of APC^Cdh1^ E3 ligase. On the other hand, the original Fucci-G1 probe had RFP fused to residues 30–120 of human Cdt1 (hCdt1(30/120)), which can serve as the direct substrate of SCF^Skp2^ E3 ligase. The original Fucci probe can be called Fucci(SA) because it monitors the balance between <u>S</u>CF^Skp2^ and <u>A</u>PC^Cdh1^ E3 ligase activities. Fucci(SA) effectively highlights the transition process from G1 phase to S phase (Figure 1A).

**Figure 1.**
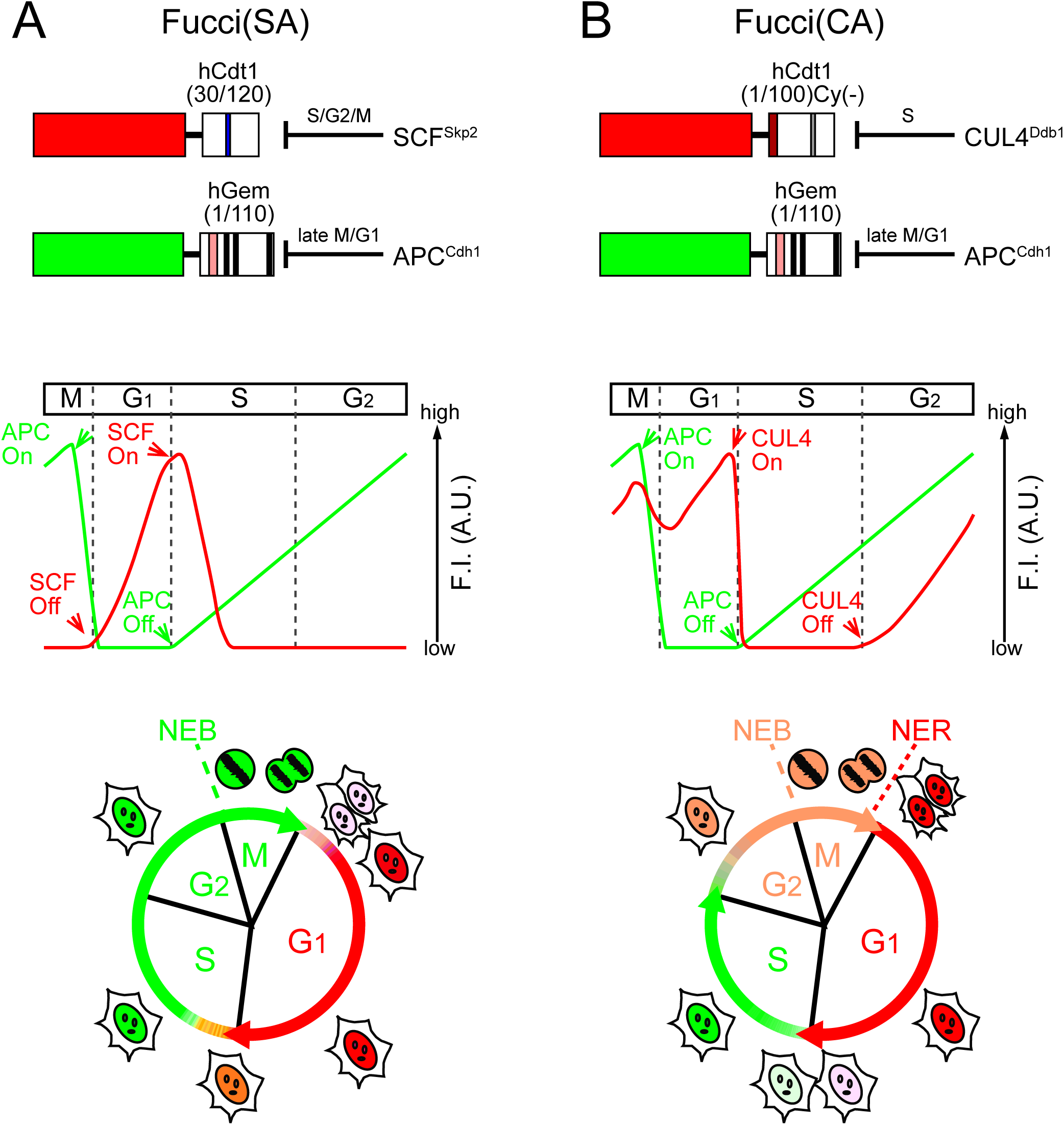
Fucci Probes with Different Ubiquitination Domains of Human Cdt1. (A) Fucci(SA) consists of an <u>S</u>CF^Skp2^-sensitive hCdt1-based probe and an <u>A</u>PC^Cdh1^-sensitive hGem-based probe. Fucci(SA) corresponds to the original Fucci. A blue box in hCdt1(30/120) indicates the Cy motif. (B) Fucci(CA) consists of a <u>C</u>UL4^Ddb1^-sensitive hCdt1-based probe and an <u>A</u>PC^Cdh1^-sensitive hGem-based probe. The dark red box and the gray box in hCdt1(1/100)Cy(-) indicates the PIP box and Cy(-): mutated Cy motif, respectively. (A, B) Domain structures (top) and cell-cycle phasing capabilities (bottom) are shown, assuming that the hCdt1- and hGem-based domains are fused to red- and green-emitting FPs. A theoretical temporal profile of the fluorescence intensity (F.I.) is shown below each domain structure. SCF, SCF^Skp2^; CUL4, CUL4^Ddb1^; APC, APC^Cdh1^. Pink and black boxes in hGem(1/110) indicate the destruction box and nuclear localization signal, respectively. NEB: nuclear envelope breakdown. NER: re-formation of the nuclear envelope.

However, visualizing cell-cycle transitions other than G1/S is just as important. Thus, we next engineered the hCdt1-based RFP-containing probe to make it sensitive to <u>C</u>UL4^Ddb1^ instead of <u>S</u>CF^Skp2^. By combining the resultant probe with the hGem(1/110)-containing green or yellow probe sensitive to <u>A</u>PC^Cdh1^, we developed Fucci(CA), which monitors the balance between <u>C</u>UL4^Ddb1^ and <u>A</u>PC^Cdh1^ (Sakaue-Sawano et al., 2017). Fucci(CA) distinguishes clearly interphase boundaries between G1, S, and G2 phases (Figure 1B).

Fucci probes have been diversifying in color in the past decade. Initially, Fucci employed mAG (monomeric Azami Green) and mKO2 (monomeric Kusabira Orange2) (Sakaue-Sawano et al., 2008). Then, the 2^nd^ generation, including Fucci(SA)2 and Fucci(CA)2, used mVenus and mCherry (Sakaue-Sawano et al., 2011; Mort et al., 2014: Sakaue-Sawano et al., 2017). Recently, far-red or near-infrared FPs have been substituted to generate Fucci variants for intravital deep imaging (Rodriguez et al., 2016; Shchebakova et al., 2016). However, although Fucci technology has become a standard method for cell-cycle analysis in academe (Zielke and Edgar, 2015; Greenwald et al., 2018), its spread to industry has been limited due to FP-license-related problems.

In the present study, we replaced the mVenus/mCherry pair in both Fucci(SA)2 and Fucci(CA)2 probes with a new GFP/RFP pair of our own cloning and engineering. We developed practically useful GFP and RFP named h2-3 and AzaleaB5, respectively, from corals. Then, we examined if h2-3 and AzaleaB5 could be substituted to develop new Fucci probes that would outperform the conventional ones and/or become more widely accepted than before.

## Results and Discussion

We screened approximately 100,000 bacterial colonies containing a cDNA library prepared from *Montipora monasteriata* (Figure 2A) for fluorescence. One clone was selected that appeared to encode an RFP, and temporarily referred to as 1-41. Based on an amino acid sequence alignment (Figure 2B), 1-41 was supposed to have a similar β-can fold to other common FPs. The closest homologue was pporRFP, an RFP cloned from *Porites porites* (Poritiina, Poritidae) (Alieva et al., 2008), which shared 80.8% identity. Transformation of the cDNA into *Escherichia coli* generated dim red fluorescent colonies. The addition of a histidine6 tag at the N-terminus of the protein allowed purification by metal affinity chromatography for spectroscopic and biochemical characterizations. The absorption spectrum of 1-41 at pH 7.4 displayed a major peak at 573 nm (Figure 2C) and a slight shoulder at 537 nm; a small peak at 503 nm was indicative of a green-emitting byproduct (Miyawaki et al., 2012). Excitation at around 540 nm produced weak fluorescence peaking at 592 nm (Figure 2D). Pseudo-native gel electrophoresis analysis revealed that 1-41 formed a highly multimeric complex (Figure 3).

**Figure 2.**
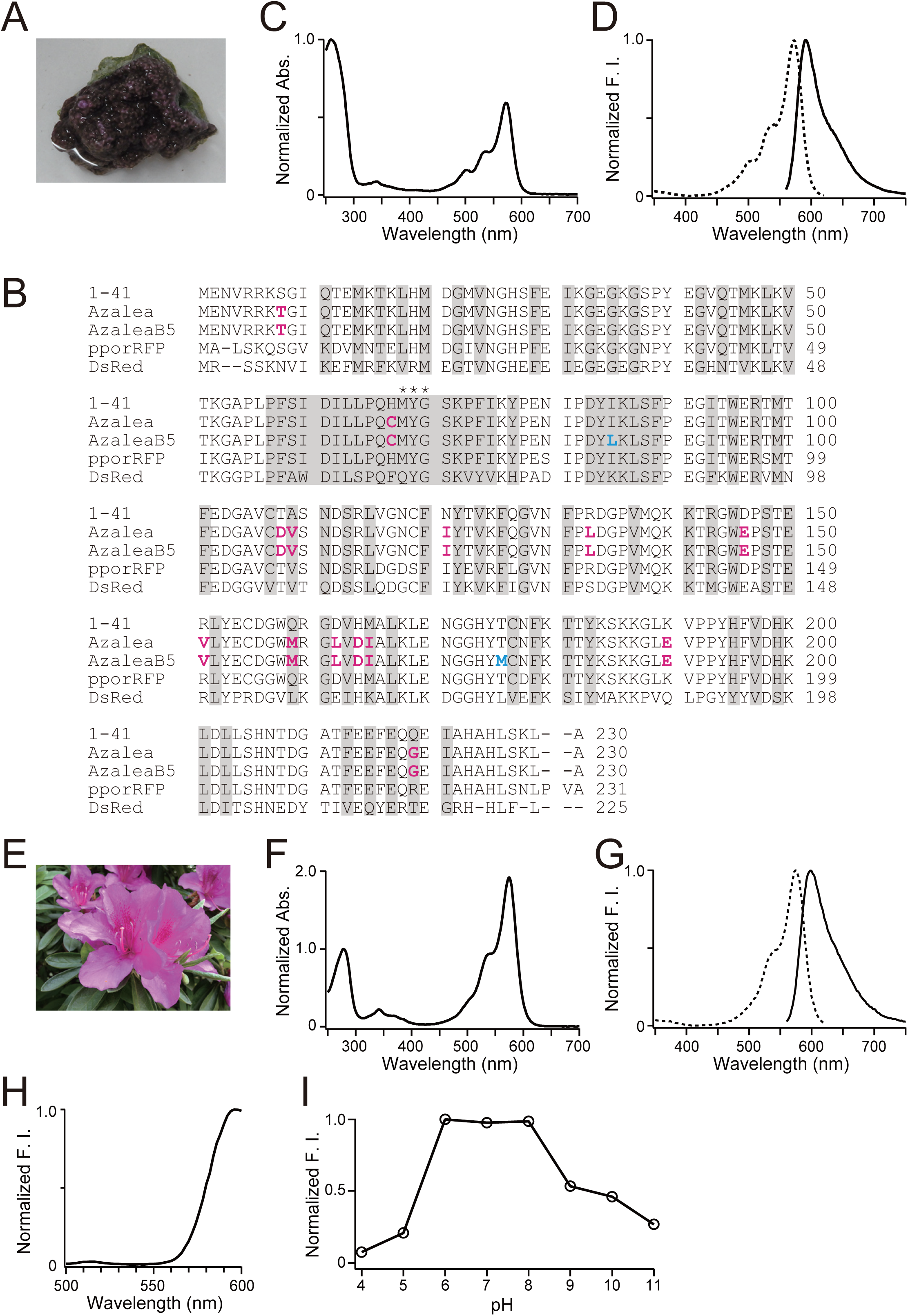
Molecular and Spectroscopic Characterizations of AzaleaB5. (A) *Montipora monasteriata*. (B) Amino acid sequence (single-letter code) alignments of 1-41, Azalea, AzaleaB5, pporRFP, and DsRed. Residues whose side chains form the interior of the β-barrel are shaded. Residues responsible for chromophore synthesis are indicated by asterisks. In the sequences of Azalea and AzaleaB5, the substituted amino acids in comparison with 1-41 are indicated in magenta. In the sequence of AzaleaB5, the substituted amino acids in comparison with Azalea are indicated in cyan. (C) Absorption spectrum of 1-41. The spectrum is normalized by the peak at 260 nm. (D) Normalized excitation (dotted line) and emission (solid line) spectra of 1-41. F.I., fluorescence intensity. (E) Azalea. (F) Absorption spectrum of AzaleaB5. The spectrum is normalized by the peak at 280 nm. (G) Normalized excitation (dotted line) and emission (solid line) spectra of AzaleaB5. F.I., fluorescence intensity. (H) Emission spectrum of AzaleaB5 with excitation at 480 nm. F.I., fluorescence intensity. (I) pH dependence of the fluorescence of AzaleaB5. F.I., fluorescence intensity.

**Figure 3.**
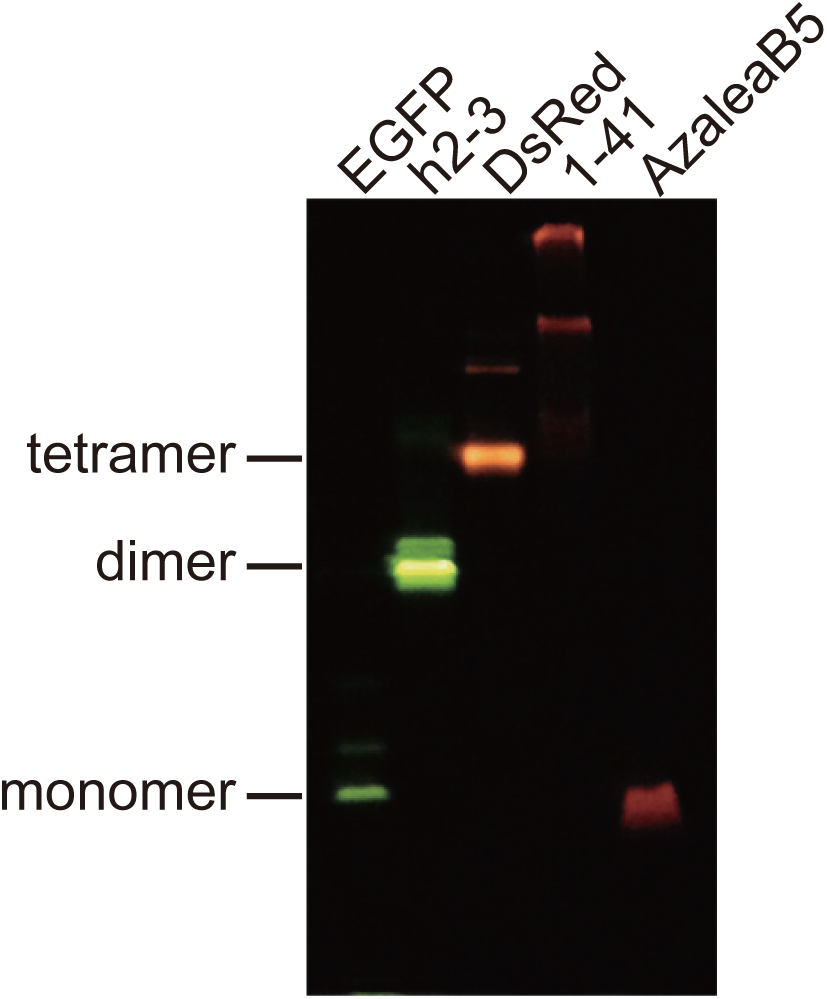
Pseudo-Native Gel Electrophoresis Analysis. EGFP and DsRed were used as size markers (monomer and tetramer, respectively). The gel was illuminated with UV light (365 nm) and imaged using a color CCD camera.

We adopted semi-random mutagenesis to transform 1-41 into a useful RFP. We performed site-directed mutagenesis to break the multimeric structure, followed by random mutagenesis to rescue the red fluorescence (Campbell et al., 2002; Ando et al., 2004). We first introduced 14 mutations: S8T, H67C, T108D, A109V, N121I, R133L, D146E, R151V, Q159M, D162L, H164D, M165I, K190E, and Q219G into #1-41 (Figure 2B). Among them, T108D was introduced into the AB interface, and D146E, R151V, D162L, and H164D were introduced into the AC interface. We found that M165I was effective in increasing the photostability of the red fluorescence. The resultant RFP was practically bright and named “Azalea” after Wako City’s designated flower (Figure 2E). Next, we used Azalea as the parental FP to develop several better mutants. One of them was AzaleaB5, which was generated by adding I85L and T176M into Azalea. Apparently, these two mutations further improved both brightness and folding efficiency. We also introduced silent base changes to optimize the coding sequence based on human codon-usage preferences. The absorption spectrum of AzaleaB5 at pH 7.4 displayed a major absorption maximum at 574 nm (ε = 104,000 M^-1^·cm^-1^) with a slight shoulder around 542 nm (Figure 2F). Excitation and emission spectra were analyzed to characterize the red-emitting component (Figure 2G); the fluorescence quantum yield (QY) was 0.58. The spectral characteristics of AzaleaB5 are summarized in Table 1. Excitation at 480 nm gave a negligible green emission compared with the red one (Figure 2H), indicating that AzaleaB5 was free from contamination by the green-emitting component. The red fluorescence was stable at pH 6–8, but decreased with increasing acidity and alkalinity (Figure 2I). Such alkaline sensitivity seemed to be unique to AzaleaB5; most conventional RFPs were stable in an alkaline as well as a neutral pH region. In pseudo-native gel electrophoresis, AzaleaB5 appeared to behave as a monomer (Figure 3).

**Table 1.**
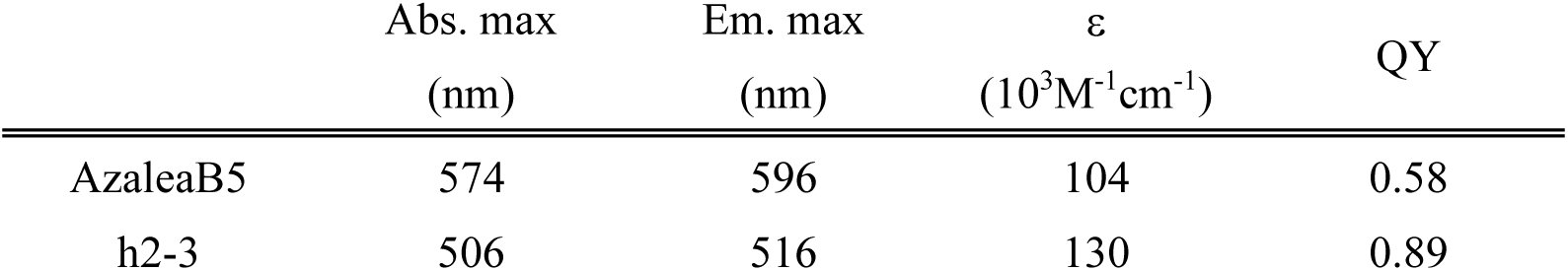
Spectral properties of AzaleaB5 and h2-3 ε: molar extinction coefficient. QY: fluorescence quantum yield.

We also cloned a cDNA that encoded a bright green-emitting FP from *Ricordea sp*. (Figure 4A). The FP was temporarily referred to as 2-3. We also generated a mutated cDNA that encoded 2-3 with human codon-usage preferences. The resultant FP was named h2-3. Sequence analysis revealed that its nearest homologue was sarcGFP from *Sarcophyton sp*. (Octocorallia, Alcyoniidae) (Figure 4B) (Alieva et al., 2008). The absorption spectrum of h2-3 at pH 7.4 displayed a major absorption maximum at 506 nm (ε = 130,000 M^-1^·cm^-1^) with a slight shoulder around 479 nm (Figure 4C). The protein exhibited an emission spectrum peaking at 516 nm (Figure 4D), which was sensitive to acidity with a p*K*_a_ of 4.6 (Figure 4E). The spectral characteristics of h2-3 are summarized in Table 1. It was shown by pseudo-native gel electrophoresis analysis that h2-3 formed an obligate dimeric complex (Figure 3).

**Figure 4.**
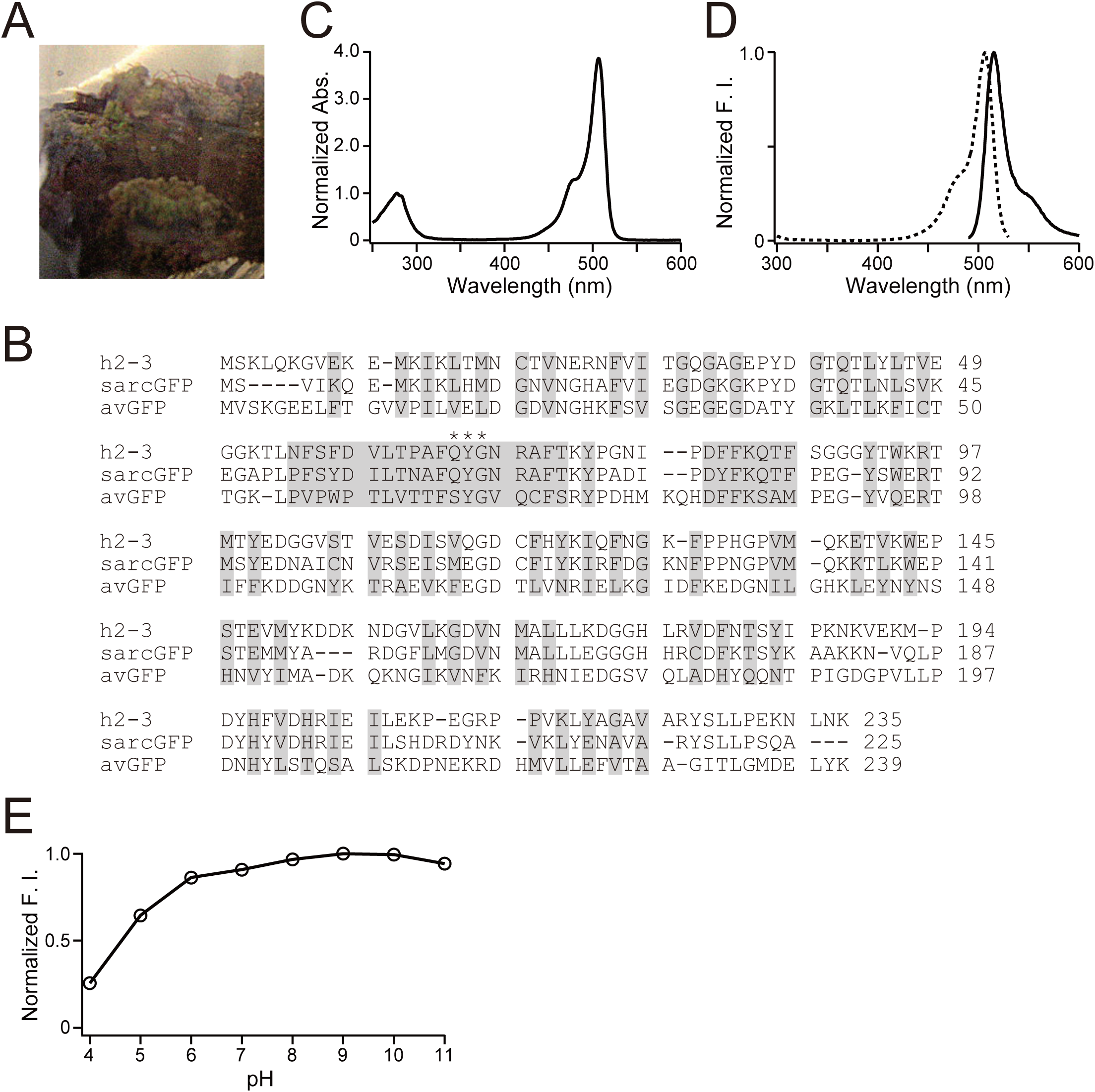
Molecular and Spectroscopic Characterizations of h2-3. (A) *Ricordea sp*. (B) Amino acid sequence (single-letter code) alignments of h2-3, sarcGFP, and *Aequorea victoria* GFP (avGFP). Residues whose side chains form the interior of the β-barrel are shaded. Residues responsible for chromophore synthesis are indicated by asterisks. (C) Absorption spectrum of h2-3. The spectrum is normalized by the peak at 280 nm. (D) Normalized excitation (dotted line) and emission (solid line) spectra of h2-3. F.I., fluorescence intensity. (E) pH dependence of the fluorescence of h2-3. F.I., fluorescence intensity.

Conventional Fucci2 (Sakaue-Sawano et al., 2011; Mort et al., 2014) is identical to Fucci(SA)2, which is composed of mCherry-hCdt1(30/120) and mVenus-hGem(1/110). On the other hand, Fucci(CA)2 is composed of mCherry-hCdt1(1/100)Cy(-) and mVenus-hGem(1/110). As Fucci(SA)2 and Fucci(CA)2 shared mVenus-hGem(1/110), we first manipulated this Geminin-based probe. Interestingly, hGem(1/110) can be fused to an FP that forms an obligate multimeric complex. For example, AmCyan, which forms an obligate tetramer, was successfully fused to hGem(1/110) to label S–G2–M-phase nuclei cyan (Nishimura et al., 2013; Sakaue-Sawano et al., 2017). Thus, we reasoned that the dimeric complex formation of h2-3 should not be a problem for fusion to hGem(1/110) to construct h2-3-hGem(1/110). Next, we constructed AzaleaB5-hCdt1(30/120) and AzaleaB5-hCdt1(1/100)Cy(-), and combined them with h2-3-hGem(1/110) to develop Fucci(SA)5 and Fucci(CA)5, respectively (Figures 5A and 5B, top). The suffix “5” indicates that AzaleaB5 and h2-3 are used for fluorescence labeling. Furthermore, it was desirable that the Cdt1-based and Geminin-based probes be concatenated via the 2A peptide and encoded by a single transgene. This tFucci (tandem Fucci) approach guaranteed the stoichiometry of the two probes, thereby enhancing the reproducibility of cell-cycle analysis data obtained from cultured cells (Sakaue-Sawano et al., 2017) and developing mouse embryos (Mort et al., 2014). Accordingly, we constructed tFucci(SA)5 and tFucci(CA)5 (Figures 5A and 5B, respectively, bottom), which were subcloned into pPBbsr2 vector. These plasmid DNAs were used to generate HeLa cells that stably expressed Fucci(SA)5 or Fucci(CA)5.

**Figure 5.**
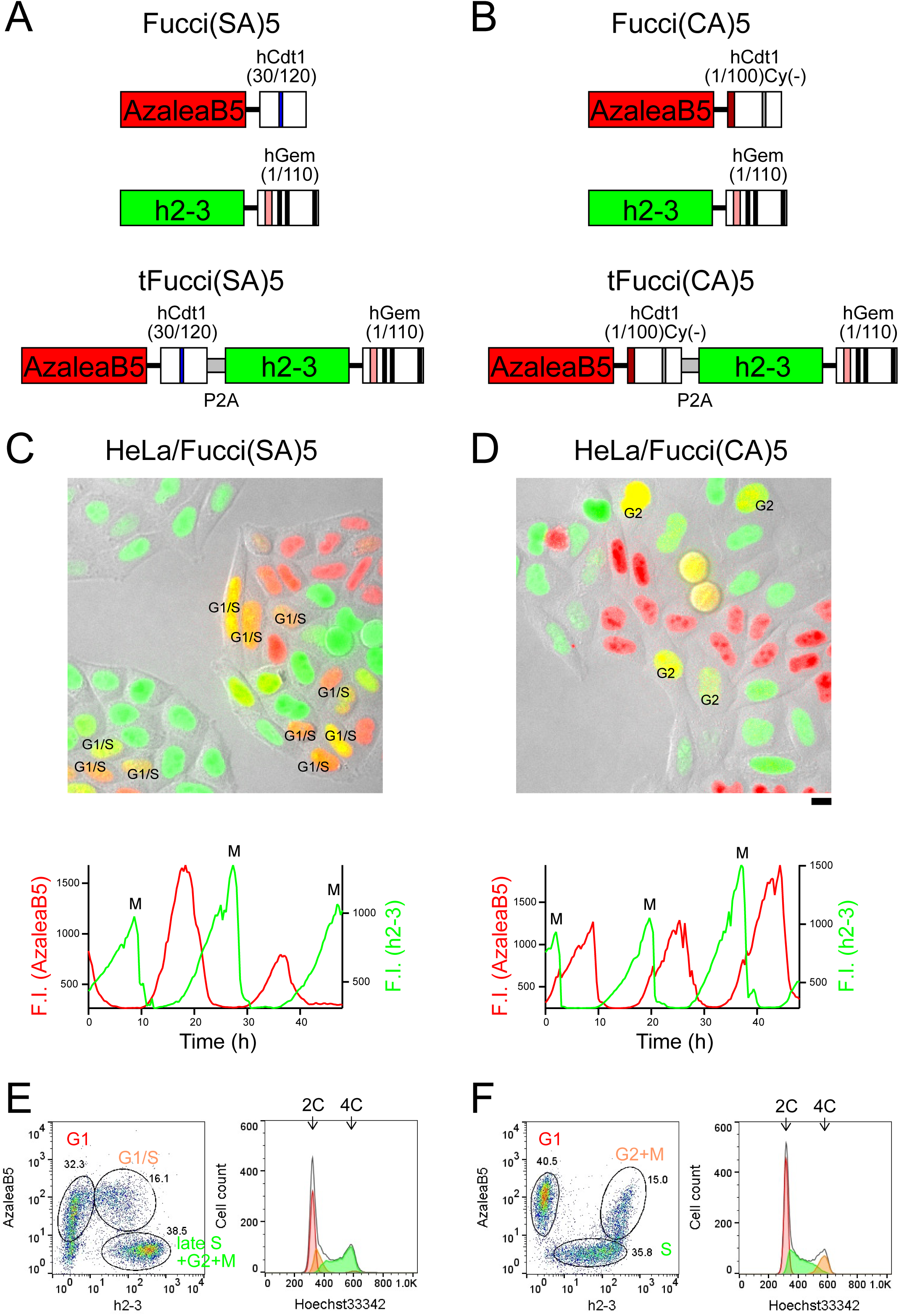
Characterization of Fucci(SA)5 and Fucci(CA)5 for Cell-Cycle Progression in HeLa Cells. (A) Fucci(SA)5 and its tandem Fucci variant, tFucci(SA)5. (B) Fucci(CA)5 and its tandem Fucci variant, tFucci(CA)5. (C, D) Time-lapse imaging of HeLa/Fucci(SA)5 (C) and HeLa/Fucci(CA)5 (D). Fucci fluorescence and DIC images were merged. These cells were in the exponentially growing phase. Images were taken every 20 min and each experiment spanned 48 h. *top*, Snapshot images. Cells in G1/S transition (C) and cells in G2 phase (D) are highlighted. *bottom*, Temporal profiles of fluorescence intensities (F.I.) of AzaleaB5 and h2-3 are indicated by red and green lines, respectively. M, mitosis. Scale bar, 10 μm. (E, F) Flow cytometry analyses of HeLa/Fucci(SA)5 (E) and HeLa/Fucci(CA)5 (F). *left*, Cells showing red [AzaleaB5(+)h2-3(-)], yellow [AzaleaB5(+)h2-3(+)], and green [AzaleaB5(-)h2-3(+)] fluorescence were gated for quantification of their DNA contents by staining with Hoechst 33342. *right*, C values denote DNA content as a multiple of the normal haploid genome.

We examined the temporal profiles of the fluorescence intensities of AzaleaB5 and h2-3 by single-cell tracking analysis under a light microscope (Figures 5C and 5D, Table 2, Movies S1 and S2). We also investigated the cell-cycle monitoring behaviors of Fucci(SA)5 and Fucci(CA)5 by population analysis (Figures 5E and 5F, respectively). After staining with Hoechst 33342 for 30 min, the cells were harvested and analyzed alive by flow cytometry (Table 2). We noted that the cells labeled with yellow fluorescence by Fucci(SA)5 and Fucci(CA)5 had DNA contents of 2–4 C and 4 C, respectively.

**Table 2.**
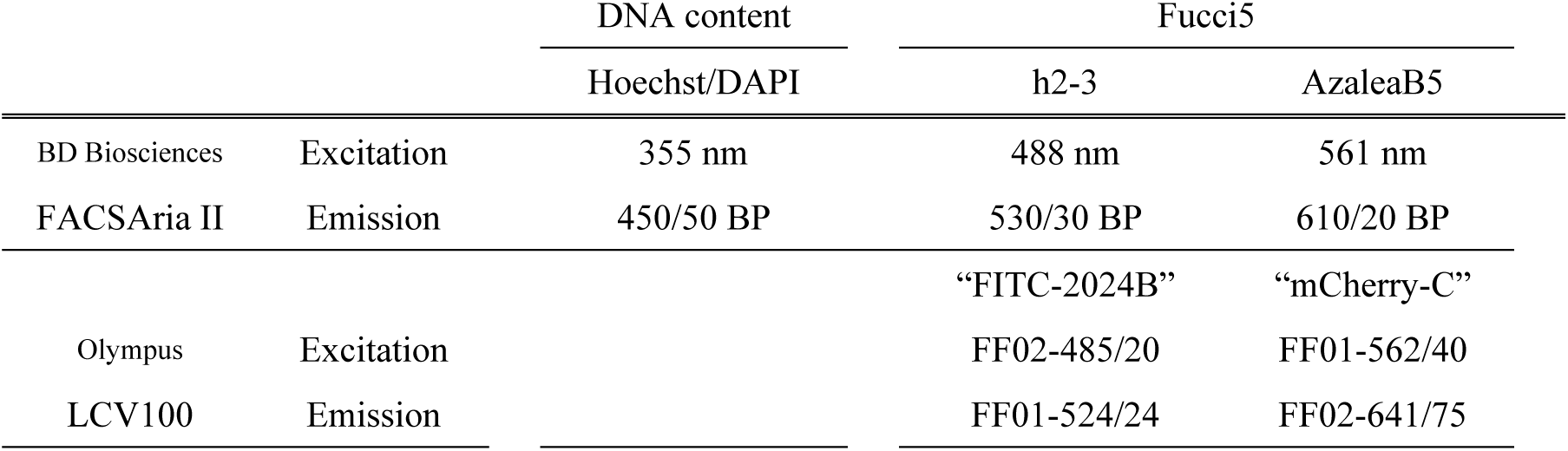
Optical components used for flow cytometry and time-lapse imaging

|  |  | DNA content | Fucci5 |  |
| --- | --- | --- | --- | --- |
|  |  | Hoechst/DAPI | h2-3 | AzaleaB5 |
| BD Biosciences | Excitation | 355 nm | 488 nm | 561 nm |
| FACSAria II | Emission | 450/50 BP | 530/30 BP | 610/20 BP |
|  |  |  | “FITC-2024B” | “mCherry-C” |
| Olympus | Excitation |  | FF02-485/20 | FF01-562/40 |
| LCV100 | Emission |  | FF01-524/24 | FF02-641/75 |

The spectral properties of AzaleaB5 and h2-3 with the optical components of confocal fluorescence microscopy for dual-color imaging are shown in Figure 6. These two FPs are excited best by common laser lines (488 and 561 nm) and their emissions are collected efficiently and specifically using a conventional multichroic mirror for the two laser lines. We have generated a human fibrosarcoma cell line (HT1080) that stably expressed tFucci(SA)5 or tFucci(CA)5. We confirmed that both HT1080/Fucci(SA)5 and HT1080/Fucci(CA)5 cells exhibited very bright nuclear labeling with green or red fluorescence. Likewise, we generated stable transformants of tFucci(SA)5 and tFucci(CA)5 using the human hepatocellular carcinoma cell line HepG2: HepG2/Fucci(SA)5 and HepG2/Fucci(CA)5, respectively. These Fucci5 cell lines will soon be used not only in the fields of basic life science and medical science but also extensively in the biotechnology industry, beginning with the drug discovery industry.

**Figure 6.**
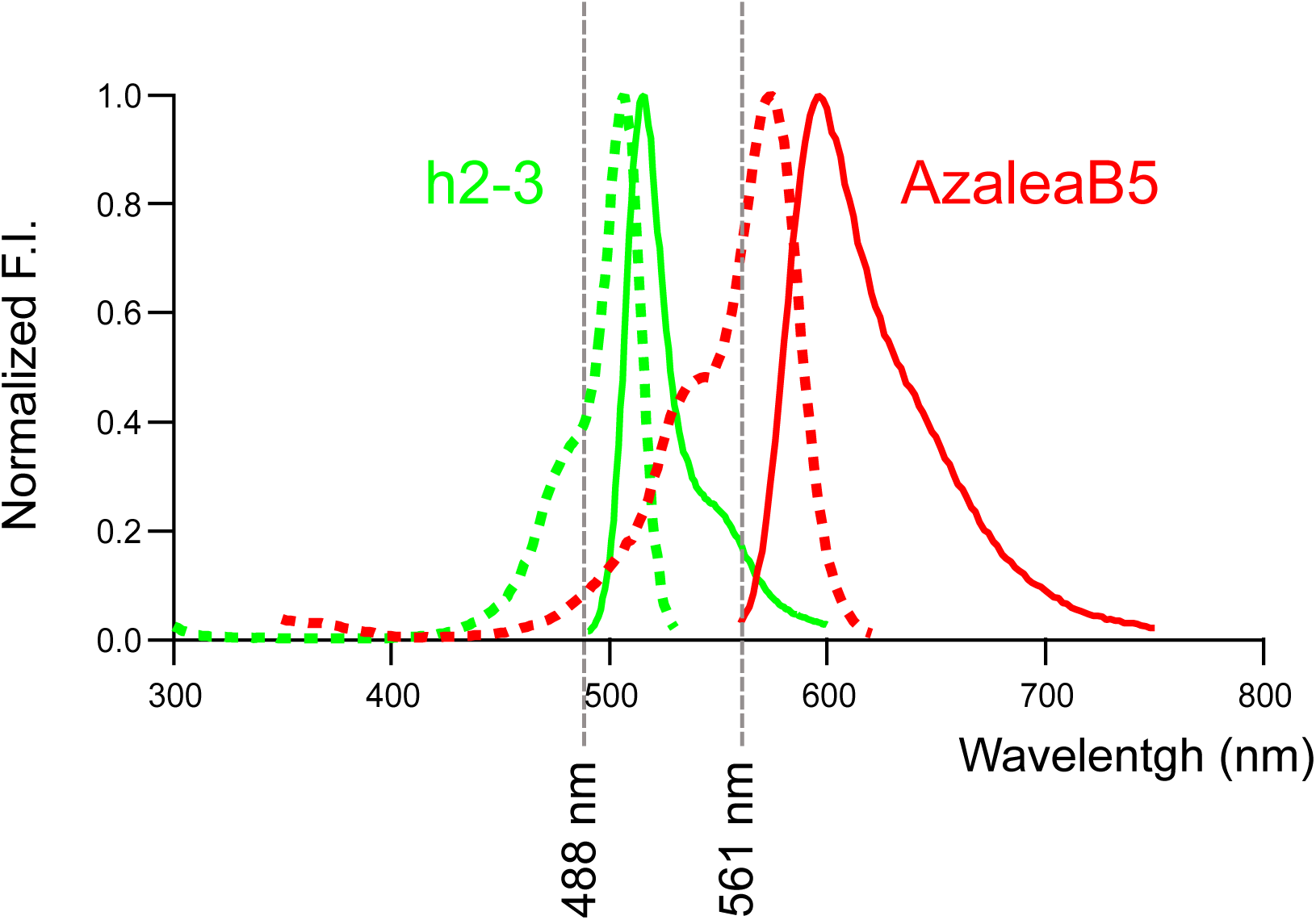
Spectral Properties of AzaleaB5 and h2-3 with Laser Wavelengths. Normalized excitation (dotted line) and emission (solid line) spectra of AzaleaB5 (red) and h2-3 (green) are shown.

## Acknowledgments

The authors thank K. Ohtawa, T. Kogure, and RIKEN CBS-Olympus Collaboration Center (BOCC) for technical assistance, J. Suzuki (RIKEN Cluster for Industry Partnerships) for valuable support, and Y. Okada (RIKEN Center for Biosystems Dynamics Research) for advice. This work was supported in part by grants from the Japan Ministry of Education, Culture, Sports, Science and Technology Grant-in-Aid for Scientific Research (B) (19H03140 to A.S-S), Scientific Research on Innovative Areas: Resonance Bio (15H05948 to A.M), and Living in Space (18H04990 to A.S-S), and the Brain Mapping by Integrated Neurotechnologies for Disease Studies (Brain/MINDS) (AMED-CREST) from Japan Agency for Medical Research and Development, AMED, and RIKEN Technology Transfer Support Fund.

## Data and materials availability

Genes: The AzaleaB5, h2-3, tFucci(SA)5, and tFucci(CA)5 genes will be available from the RIKEN BioResource Research Center (BRC) at Tsukuba (http://en.brc.riken.jp/) under a material transfer agreement with RIKEN. The accession numbers in the DDBJ/EMBL/GenBank databases are [LC085679] for AzaleaB5, [LC085680] for h2-3 (under the name of FP2-3h), [LC334437] for tFucci(SA)5, and [LC334438] for tFucci(CA)5.

Stable cell lines: HeLa/Fucci(SA)5 (clone #16) and HeLa/Fucci(CA)5 (clone #16) cells will be distributed by the RIKEN BioResource Research Center (BRC) Cell Bank (https://cell.brc.riken.jp/en/).

The information about Fucci-related materials is available in our website (http://cfds.brain.riken.jp/Fucci.html).

## Methods

### cDNA Cloning

The soft coral *Ricordia sp*. and the stony coral *Montipora monasteriata* were purchased from an aquarium shop. For each coral, whole tissue was frozen and ground down with a MultiBeads Shocker (Yasui Kikai), and total RNA was isolated by TRIzol Reagent (Thermo Fisher Scientific). mRNAs were purified using an Oligotex-dT30 <Super> (JSR). cDNA was synthesized with an *Sal*I site at the 5’ end and a *Not*I site at the 3’ end by using a SuperScript™ Plasmid System with Gateway^®^ Technology for cDNA Synthesis and Cloning (Thermo Fisher). Ligation of the cDNAs into an *Sal*I/*Not*I-cleaved pRSET-FastBac plasmid (Ando et al., 2002) produced a directional cDNA library in a prokaryotic expression vector. The libraries were transformed into the *E. coli* strain JM109 (DE3). Colonies were screened for fluorescence by using a UV illuminator (365 nm) and a LED transilluminator (green).

### Mutagenesis

Site-directed and semi-random mutations were introduced according to our protocols as described previously (Sawano and Miyawaki, 2000). Error-prone mutagenesis was based on PCR using GoTaq DNA polymerase (Promega) supplemented with 1 mM MnCl_2_.

### Protein Expression, Spectroscopy, pH titration

The cDNA of the coding region of fluorescent proteins were amplified by using primers containing 5’ *Bam*HI and 3’ *Eco*RI sites. The restricted products were cloned in-frame into the *Bam*HI/*Eco*RI site of pRSETB for bacterial expression. Proteins were expressed in *E. coli* and purified by Ni-NTA (QIAGEN). Then protein samples were desalted through a PD-10 column (GE Healthcare). *In vitro* spectroscopy was performed in 50 mM, HEPES-NaOH, pH 7.4. Absorbance spectra were acquired with a spectrophotometer (U-3310, Hitachi). Fluorescence measurements were performed using a microplate spectrophotometer (SynergyMx, BioTek). pH titration buffers used were below; 50 mM NaOAc-HOAc (pH 4.0–5.0), 50 mM KH_2_PO_4_-NaOH (pH 6.0), 50 mM HEPES-NaOH (pH 7.0–8.0), 50 mM Glycine-NaOH (pH 9.0–10.0), 50 mM Na_2_HPO_4_-NaOH (pH 11.0), 50 mM KCl-NaOH (pH 12.0). Assuming the molar extinction coefficient for denatured chromophore was 44,000 M^-1^cm^-1^, molar extinction coefficients of FPs were calculated by the ratio of matured chromophore absorbance and denatured chromophore absorbance (Shaner et al., 2004). The fluorescence quantum yields were measured by an absolute PL quantum yield spectrometer (C9920-02, Hamamatsu photonics) in 50 mM HEPES-NaOH, pH7.4.

### Pseudo-Native Gel Electrophoresis

Purified proteins were mixed with 4× sample buffer (0.2 M Tris-HCl, pH 6.8, 8% SDS, 20% 2-mercaptoethanol, 40% glycerol, 0.4% BPB) and run on a 10% polyacrylamide gel without denaturation. The gel was imaged with a digital color CCD camera under UV irradiation.

### Gene Construction

tFucci probes were constructed by concatenating the hCdt1-based probe, P2A sequence (Fang et al., 2005; Kim et al., 2011), and the hGem-based probe. We utilized a PiggyBac transposon system to generate cells that stably express tFucci probes (Aoki et al k., 2013). The mVenus-P2A-mCherry-hGem(1/110) gene in pPBbsr2 was used for construction of tFucci(SA)5. DNA fragments encoding *Bam*HI-AzaleaB5-*Eco*RV-*Not*I-*Xho*I and *Xho*I-hCdt1(30/120)-P2A-*Eco*RI were amplified using primers, and digested products were substituted for the *Bam*HI-mVenus-P2A-*Eco*RI gene in mVenus-P2A-mCherry-hGem(1/110) in pPBbsr2 vector to produce AzaleaB5-hCdt1(30/120)-P2A-mCherry-hGem(1/110) in pPBbsr2. Then, DNA fragments encoding *Eco*RI-h2-3-*Not*I and *Not*I-hGem(1/110)-*Hpa*I were amplified using primers, and digested products were substituted for *Eco*RI-mCherry-hGem(1/110)-*Hpa*I gene in AzaleaB5-hCdt1(30/120)-P2A-mCherry-hGem(1/110) in pPBbsr2 vector. The final product AzaleaB5-hCdt1(30/120)-P2A-h2-3-hGem(1/110) was referred to as tFucci(SA)5 and the sequence has been deposited in the DDBJ/EMBL/GenBank data-base [LC334437]. Likewise, DNA fragments encoding *Bam*HI-AzaleaB5-*Xho*I and *Xho*I-hCdt1(1/100)Cy(-)-*Age*I were amplified using primers, and digested products were substituted for *Bam*HI-AzaleaB5-hCdt1(30/120)-*Age*I gene in AzaleaB5-hCdt1(30/120)-P2A-h2-3-hGem(1/110)in pPBbsr2 vector. The final product AzaleaB5-hCdt1(1/100)Cy(-)-P2A-h2-3-hGem(1/110) was referred to as tFucci(CA)5 and the sequence has been deposited in the DDBJ/EMBL/GenBank data-base [LC334438].

### Cell Culture

HeLa cells (a subclone of HeLa.S3) were grown in Dulbecco’s modified Eagle’s medium (DMEM) (FUJIFILM Wako Pure Chemical Cooperation) supplemented with 10% fetal bovine serum (FBS) and penicillin/streptomycin. HeLa.S3 has been characterized to proliferate relatively fast with a doubling time of 15–18 hours (Sakaue-Sawano et al., 2008).

### Establishment of Stable Cell Lines

For generation of HeLa cell lines stably expressing tFucci probes, the PiggyBac transposon system was employed (Aoki et al., 2013). The pPBbsr-based tFucci probes and pCMV-mPBase (neo-) encoding the *piggyBac* transposase were cotransfected into HeLa cells using PEI (Polyethylenimine) at a ratio of 3:1. Transfected cells were selected with blasticidin S (InvivoGen) (50 μg/ml for 3 days and subsequently 10 μg/ml for 7–10 days). tFucci-expressing singe cell clones were further isolated by limited dilution.

### Flow Cytometry

Hoechst 33342 solution (56 μl of 1 mg/ml stock) (DOJINDO, Kumamoto, Japan) was added to a 10-cm dish containing HeLa/Fucci cells. After incubation for 30 min, cells were harvested and analyzed using a FACSAria II (BD Bioscience, San Jose, CA). h2-3 was excited by a 488-nm laser line (laser diode) and its emission was collected through 530/30BP; AzaleaB5 was excited by a 561-nm laser line and its emission was collected through 610/20 BP. Hoechst 33342 was excited by a UV Laser at 355 nm, and its emission was collected through 450/50 BP. The data were analyzed using FlowJo software (Tree Star). See Table 2 for details.

### Long-Term Time-Lapse Imaging

Cells were grown on 35-mm glass-bottom dishes in phenol red-free DMEM containing 10% FBS. Cells were subjected to long-term, time-lapse imaging using a computer-assisted fluorescence microscope (Olympus, LCV100) equipped with an objective lens (Olympus, UAPO 40×/340 N.A. = 0.90), a halogen lamp, a red LED (620 nm), a CMOS camera (Hamamatsu Photonics, ORCA-Flash4.0), differential interference contrast (DIC) optical components, and interference filters. The halogen lamp was used with BrightLine^®^ single-band filter set (Semrock): “FITC-2024B” for observing the h2-3 fluorescence, and “mCherry-C” for observing the AxaleaB5 fluorescence. For DIC imaging, the red LED was used with a filter cube containing an analyzer. Image acquisition and analysis were performed using MetaMorph 6.37 and 7.10 software (Molecular Devices), respectively. See Table 2 for details.

### Manual Cell Tracking

Image processing was performed manually using the “Journal” functions implemented in MetaMorph (Molecular Devices). First, fluorescence images of AzaleaB5 and h2-3 were merged. In addition, DIC images acquired at slightly different focal planes were merged for delineating individual cell nuclei. This morphology observation was particularly useful for marking mitotic events. Time sequence data of tracked cells are saved in “TrackRef” files. The mean fluorescence intensities of tracked nuclei were calculated using the “Region measurements” function.

---

Movie S1 HeLa/Fucci(SA)5 cells were grown on a glass-bottom dish, and time-lapse imaging was performed using an LCV100 microscope. Images were acquired every 17 min. Total imaging time = 48 hr.

Movie S2 HeLa/Fucci(CA)5 cells were grown on a glass-bottom dish, and time-lapse imaging was performed using an LCV100 microscope. Images were acquired every 17 min. Total imaging time = 48 hr.

## References

Alieva, N.O., Konzen, K.A., Field, S.F., Maleshkevitch, E.A., Hunt, M.E., Belrran-Ramirez, V., Miller, D.J., Wiedenmann, J., Salih, A., and Matz, M.V (2008). Diversity and evolution of coral fluorescent proteins. PLoS ONE 3, e2680.

Ando, R., Hama, H., Yamamoto-Hino, M., Mizuno, H., and Miyawaki, A. (2002). An optical marker based on the UV-induced green-to-red photoconversion of a fluorescent protein. Proc. Natl. Acad. Sci. USA 99, 12651–12656.

Ando, R., Mizuno, H., and Miyawaki, A. (2004). Regulated fast nucleocytoplasmic shuttling observed by reversible protein highlighting. Science 306, 1370–1373.

Aoki, K., Kumagai, Y., Sakurai, A., Komatsu, N., Fujita, Y., Shionyu, C., and Matsuda, M. (2013). Stochastic ERK activation induced by noise and cell-to-cell propagation regulates cell density-dependent proliferation. Mol. Cell 52, 529–540.

Campbell, R.E., Tour, O., Palmer, A.E., Steinbach, P.A., Baird, G.S., Zacharias, D.A., Tsien, R.Y. (2002). A monomeric red fluorescent protein. Proc. Natl. Acad. Sci. USA 99, 7877–7882.

Fang, J., Qian, J.J., Yi, S., Harding, T.C., Tu, G.H., VanRoey, M., and Jooss, K. (2005). Stable antibody expression at therapeutic levels using the 2A peptide. Nat. Biotechnol. 23, 584–590.

Greenwald, E.C., Mehta, S., and Zhang, J. (2018). Genetically encoded fluorescent biosensors illuminate the spatiotemporal regulation of signaling networks. Chem. Rev. 118, 11707–11794.

Kim, J.H., Lee, S.R., Li, L.H., Park, H.J., Park, J.H., Lee, K.Y., Kim, M.K., Shin, B.A., and Choi, S.Y. (2011). High cleavage efficiency of a 2A peptide derived from porcine teschovirus-1 in human cell lines, zebrafish and mice. PLoS One 6, e18556.

Miyawaki, A.M., Shcherbakova, D.M., and Verkhusha, V.V. (2012). Red fluorescent proteins: chromophore formation and cellular applications. Curr. Opin. Struct. Biol. 22, 679–688.

Mort, R.L., Ford, M. J., Sakaue-Sawano, A., Lindstrom, N.O., Casadio, A., Douglas, A.T., Keighren, M.A., Hohenstein, P., Miyawaki, A., and Jackson, I.J. (2014). Fucci2a: a bicistronic cell cycle reporter that allows Cre mediated tissue specific expression in mice. Cell Cycle 13, 2681–2696.

Nishimura, K., Oki, T., Kitaura, J., Kuninaka, S., Saya, H., Sakaue-Sawano, A., Miyawaki, A., and Kitamura, T. (2013). APC^Cdh1^ targets MgcRacGAP for destruction in the late M phase. PLoS One. 8, e63001, doi: 10.1371/journal.pone.0063001.

Rodriguez, E.A., Tran, G.N., Gross, L.A., Crisp, J.L., Shu, X., Lin, J.Y., and Tsien, R.Y. (2016). A far-red fluorescent protein evolved from a cyanobacterial phycobiliprotein. Nat. Methods 13, 763–769.

Rodriguez, E.A., Campbell, R.E., Lin, J.Y., Lin, M.Z., Miyawaki, A., Palmer, A.E., Shu, X., Zhang, J., and Tsien, R.Y. (2017). The growing and glowing toolbox of fluorescent and photoactive proteins. Trends. Biochem. Sci. 42, 111–129.

Sakaue-Sawano, A., Kurokawa, H., Morimura, T., Hanyu, A., Hama, H., Osawa, H., Kashiwagi, S., Fukami, K., Miyata, T., Miyoshi, H., et al. (2008). Visualizing spatiotemporal dynamics of multicellular cell-cycle progression. Cell 132, 487–498.

Sakaue-Sawano, A., Kobayashi, T., Ohtawa, K., and Miyawaki, A. (2011). Drug-induced cell cycle modulation leading to cell-cycle arrest, nuclear mis-segregation, or endoreplication. BMC Cell Biol. 13, 12:2.

Sakaue-Sawano, A., Yo, M., Komatsu, N., Hiratsuka, T., Kogure, T., Hoshida, T., Goshima, N., Matsuda, M., Miyoshi H., and Miyawaki, A. (2017). Genetically Encoded Tools for Optical Dissection of the Mammalian Cell Cycle. Mol Cell 68, 626–640.

Sawano, A., and Miyawaki, A. (2000). Directed evolution of green fluorescent protein by a new versatile PCR strategy for site-directed and semi-random mutagenesis. Nucleic Acids Res. 28, E78.

Shaner, N.C., Campbell, R.E., Steinbach, P.A., Giepmans, B.N., Palmer, A.E., and Tsien, R.Y. (2004). Improved monomeric red, orange and yellow fluorescent proteins derived from Discosoma sp. Red fluorescent protein. Nat. Biotechnol. 22, 1567–1572.

Shcherbakova, D.M., Baloban, M., Emelyanov, A.V., Brenowitz, M., Guo, P., and Verkhusha, V.V. (2016). Bright monomeric near-infrared fluorescent proteins as tags and biosensors for multiscale imaging. Nat. Commun. 7, 12405.

Zielke, N., and Edgar, B.A. (2015). FUCCI sensors: powerful new tools for analysis of cell proliferation. Wiley Interdiscip. Rev. Dev. Biol. 4, 469–487.

